# Role of mitochondrial genetic interactions in determining adaptation to high altitude in human population around the globe

**DOI:** 10.1101/2021.06.21.449348

**Authors:** Rahul K Verma, Kalyakulina Alena, Ankit Mishra, Mikhail Ivanchenko, Sarika Jalan

## Abstract

Physiological and haplogroup studies performed to understand high-altitude adaptation in humans are limited to individual genes and polymorphic sites. Due to stochastic evolutionary forces, the frequency of a polymorphism is affected by changes in the frequency of a near-by polymorphism on the same DNA sample making them connected in terms of evolution. Here, first we provide a method to model these mitochondrial polymorphisms as “co-mutation networks” for three high-altitude populations, Tibetan, Ethiopian and Andean. Then, by transforming these co-mutation networks into weighted and undirected gene-gene interaction (GGI) networks, we were able to identify functionally enriched genetic interactions of *CYB* and *CO3* genes in Tibetan and Andean populations, while NADH dehydrogenase genes in Ethiopian population playing a significant role in high altitude adaptation. These co-mutation based genetic networks provide insights into the role of different set of genes in high-altitude adaptation human sub-populations.

## 1. Introduction

Paramount success of network science banks heavily on influence of pair-wise interactions among the constituents of the corresponding complex systems on the evolution and performance of the systems. The current work provides a model to capture mitochondrial genomic variations by considering them as co-mutation networks, and analyse the role of polymorphic cohorts in convergent evolution for high altitude human population. The term ‘convergent evolution’ is defined as development of the same or similar phenotypic adaptations under a similar external environmental condition as a consequence of natural selection. We consider three high altitudes geographical regions, namely, (i) Andean Altiplano, (ii) Qinghai–Tibetan Plateau and (iii) the Ethiopian Highlands to understand mechanisms behind the adaptation of human life for high altitudes. It is now believed that about 25% of the mtDNA protein sequence variations [1–3], 10 to 20% of the tRNA variations, and at least few of the rRNA variations have altered the mitochondrial coupling efficiency [4], thereby leading to an increase in the heat production at the expense of ATP production eventually permitting humans to adapt to the colder climates. However, association and mutual role of polymorphisms in other regions of mtDNA are subject to investigation in establishing a conclusive role. Complex systems have been extensively studied through networks[5] to infer role of proteins through protein-protein interaction (PPI) networks[6], to understand biological functions of neurons in various developmental stages of *C. elegans* by analyzing the underlying protein-protein interactions [7], to identify crucial structural patterns in diabetes mellitus II [9], and to understand genes responsible for evolution of specific characteristics of human sub-population [10]. Networks allow modeling of real-world complex systems by a very simple yet effective framework which consists of nodes and edges. Analysing the structural and other features of the underlying network properties based on the information available of the edges (interactions) could reveal many system level properties of corresponding complex systems [11].

Indigenous human populations residing on the Tibetan, Ethiopian and Andean Plateaus are believed to be descendants of colonizers who arrived at most ~25,000, 16000 and 11,000 years ago, respectively [12]. The three high-altitude populations currently can be viewed as a outcome of separate replications of a natural experiment in which an ancestral founding population moved from low to high altitude, and its descendants have been exposed for millennia to the opportunity for natural selection to improve their functions under high-altitude hypoxia. However, the three populations are believed to be evolved differently at the physiological levels [13]. Less oxygen in the inhaled air due to high altitude results in less oxygen to diffuse into the bloodstream flowing to the cells for oxygen-requiring energy-producing metabolism in the mitochondria. Humans do not store oxygen due to its rapid and destructive reactions with other molecules. Therefore, oxygen must be supplied without interruption to the mitochondria and to the ⩾1,000 oxygen-requiring enzymatic reactions in various cells and tissues [14]. Based on factors contributing to arterial oxygen content like hemoglobin content, oxygen saturation, hemoglobin affinity, etc., there are abundant evidences of the Andean-Tibetan difference in high-altitude adaptation while investigation of the Ethiopian population lies at its nascent stage [13]. There exist certain polymorphisms belonging to genes ND3 and CYTB which are believed to be associated with high-altitude adaptation in Tajiks population native to China [15]. It has been observed that genetic variations rarely impart their effect at phenotypic levels individually but as a cohort of multiple interactions together [16]. Single nucleotide polymorphisms (SNPs) have been displayed to have a small effect on the heritability of few complex diseases [17]. Additionally, various studies have indicated that interactions of SNPs [18–20] or mutations in proteins [21] are one of the key factors in manifestation of such complex diseases. To select a particular cohort of the variations and their interactions responsible for manifestation of complex phenotypes, various computational methods have been developed and implemented such as principal component analysis to evaluate SNP correlations [22], integrative scoring system based on their deleterious effects [23], and Pareto-optimal approach [24]. There exist other approaches based on pair-wise interaction such that two variations significantly interacting through logic regression [25], predictive rule inference [26], and shrunken methodology [27]. Networks of variable sites were used to identify angiogenesis genes associated with breast cancer [28], time-dependent weight dynamics in chickens [29], feed efficiency in duroc and landrace pigs [30], altitude-dependent interactions of mitochondrial genes in Asian population [31], etc.

Here, in this work we first constructed the co-mutation network by selecting significantly interacting variations of the mitochondrial genome, thereafter, a gene-gene interaction (GGI) network was constructed from the corresponding co-mutation networks and functional enrichment analysis was performed based on significantly interacting gene-sets. These networks were found to follow the smallworld behaviour with high modularity. The weak ties which are nodes with low degree and high betweenness centrality were found only in the Tibetan network and acting as haplogroup markers. Through GGI networks, it was found that interactions of CYB and CO3 genes played important role for high-altitude adaptation in Tibetan and Andean population while ND genes in Ethiopian population.

## 2. Methods

### 2.1 Sequence Acquisition

Complete human mitochondrial genome sequences were downloaded from the Human mitochondrial Database (HmtDB) [32] for the Ethiopia and Andes regions which are situated ~3000m, and ~3500m above sea level, respectively. For Andes region, we have downloaded sequences from Peru region since it inhabits indigenous Andean (Aymara and Quechua) population. Tibetan sequences (~4000m) were downloaded from the GenBank. All the sequences were aligned globally and mapped with a master sequence rCRS (revised Cambridge Reference Sequence)[33].

### 2.2 Construction of Co-mutation Network and Gene-Gene Interaction (GGI) network

We constructed two types of networks, first, the co-mutation network where nodes are variable sites, and second, the weighted GGI network where nodes were genes (Fig 1) for each of the high-altitude population.

**Fig. 1.**
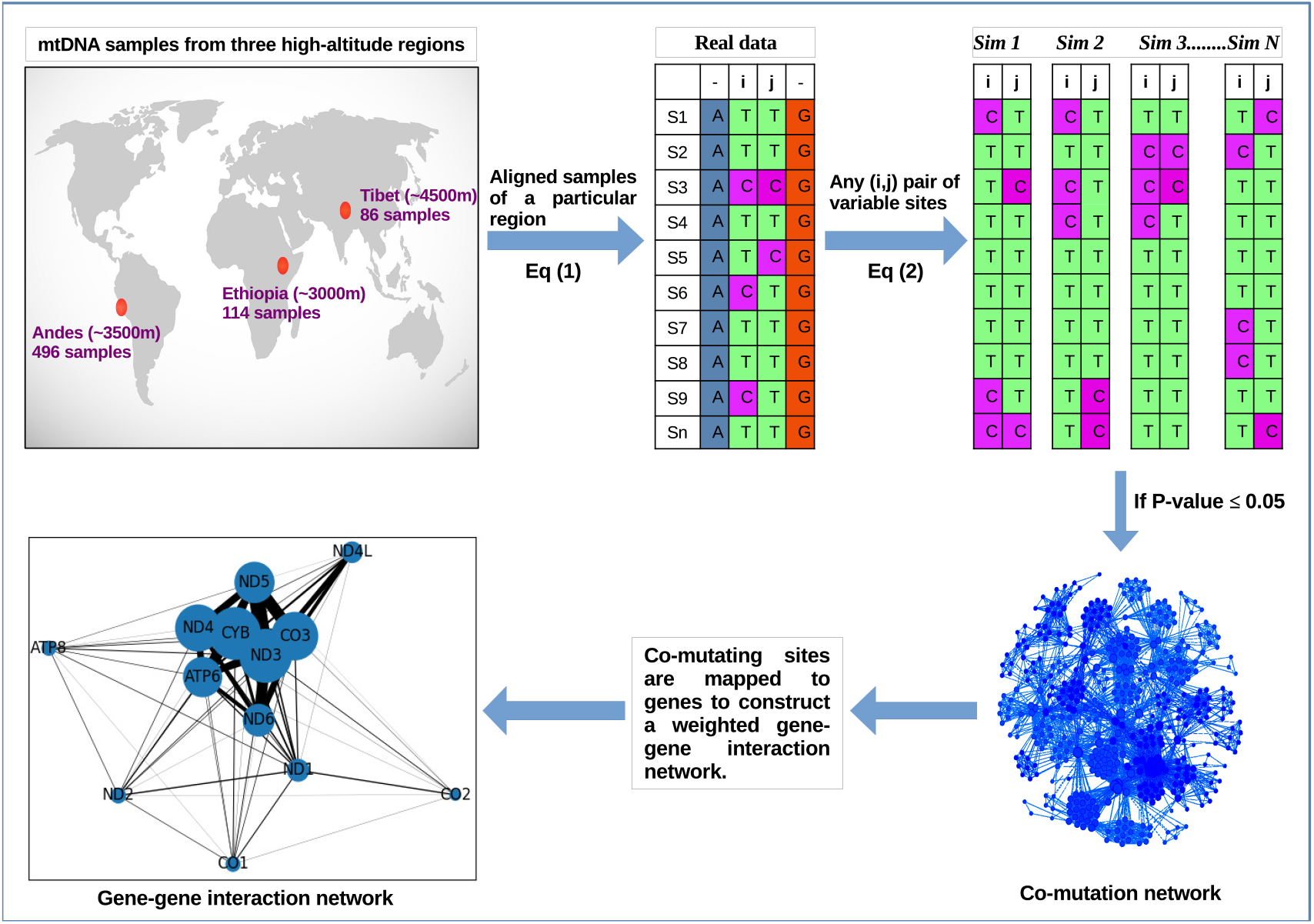
Construction of Co-mutation network and Gene-Gene Interaction (GGI) network for each high-altitude region as explained in *Methods section*.

#### Step 1 (Co-mutation Network)

Any position having more than one allele within the samples is considered as a variable site. The variable sites were extracted from the aligned sequences for each region separately. For genomic equality, ambiguous nucleotides such as X, M, Y, etc were replaced with ‘N’ for all the sequences and triallelic sites were not considered.

#### Step 2

To construct network for each high-altitude region, a node was represented by the position of a variable site and the edge was represented by co-mutation frequency between a pair of the nodes. We defined co-mutation frequency for a pair of nodes, *C_ij_*, as

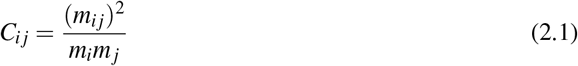

where (*m_ij_*) represents number of times the minor alleles occur together (defined as co-mutation pair) at *i^th^* and *j^th^* positions, *m_i_* and *m_j_* indicate total number of times the minor allele occurs at *i^th^* and *j^th^* position, respectively.

#### Step 3 (*p*-Value calculation)

For checking the significance of any co-mutation pair, we calculated the *p*-value as,

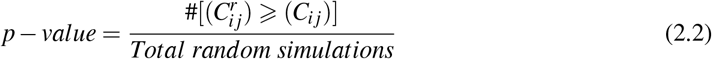

where 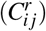 is the co-mutation frequency calculated after permuting the alleles at the *i^th^* and *j^th^* positions randomly. 10,000 random simulations were generated and *p*-value was set to ⩽ 0.05 (standard value) to consider a co-mutation pair significant.

#### Step 4 (Gene-Gene Interaction (GGI) Network)

For each co-mutation network, nodes were mapped to their corresponding genes to achieve one weighted gene-gene interaction network for each high-altitude region. Since, two or more co-mutation pairs may belong to the same gene or a gene-pair, each link is counted as many times it is found and this number was considered as weight for that genepair. For example, the co-mutation pairs (3352-7623) and (4125-8054) yield two *ND1-CO2* gene-pairs, so this pair was counted twice and so on, similarly the the co-mutation pairs (3352-3489) would yield a self-loop for *ND1* gene. The weight of a gene-pair was normalized by the sum of total length of both the genes for that gene-pair. Additionally, we have constructed GGI networks using the co-mutations of the (i) common nodes and (ii) exclusive nodes for all the three regions. For (i), we took the nodes commonly present in all the networks and scanned for their co-mutating partners, and mapped these co-mutating nodes to respective genes. Similarly, exclusive nodes for each region were used to construct the corresponding GGI network.

### 2.3 Null Model and Significant interactions

Random networks were generated corresponding to each co-mutation network to compare with the real networks by taking the same number of the nodes as in the real network, we connect them with a probability *α* such that total number of connections the random networks has is the same as of the corresponding real world;

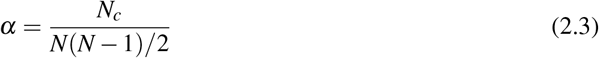

where, *N_c_* is total number of connections and *N* is total number of nodes in the real network. Further, these random networks were used to get the random GGI networks as described in *Step 4*. We compared the real GGI networks with the corresponding random GGI networks and considered only those pairs significant whose weights fell two standard deviations away from the corresponding random one.

### 2.4 Detection of Community and Role or nodes

By calculating the modularity we detected the communities using algorithm given in [34] which is implemented in Python using *community module*. It is a modularity maximization algorithm. The role of each node in the communities is determined by its within-module degree, *Z* score and the participation coefficient *P*. The within-module degree quantifies the node’s intra-modular connectivity and was calculated as the Z-score-transformed degree of centrality within the module. For a given node *i*, *Z_i_* is defined as,

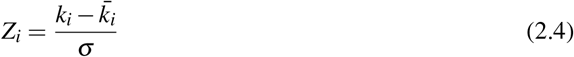

where *k_i_* is degree of the *i^th^* node in its own community, 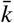 is the average of *k_i_* for all the nodes of that community, and *σ* is the standard deviation. *Z_i_* takes a high value if within the cluster, degree of *i^th^* node is high and vice versa.

Different roles of a node can also be deduced based on the number of connections the node makes with the nodes in the modules other than its own. For example, two nodes with the same Z-score will play different roles if one of them is connected to several nodes in other modules while the other is not. We define the participation coefficient *P* of node *i* as,

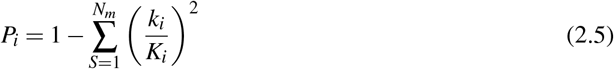

*K_i_* is the total degree of the node *i* in the whole network. *S* is the community and *N_m_* is the total number of communities. The participation coefficient of a node is therefore close to 1 if its links are uniformly distributed among all the modules and 0 if all its links are within its own module.

## 3. Results

### 3.1 Identification of Significant Interactions

We constructed the co-mutation networks for each of the high altitude human sub-population with mtDNA variants to analyse and compare the co-evolution of the variable sites for three underlined regions. The Andes region has the highest number of the samples among all the three regions (Table 1). More number of the samples rendered more number of interactions to be statistically significant which is observed in comparison of number of connections (*L*) at *C_ij_* > 0 between the Andes and Ethiopia. Both the regions are having almost equal number of connections before applying the threshold. When significant pairs (with *p*-value ⩽ 0.05) were considered, the number of connections were decreased by ~23% in Andes, ~49% in Ethiopia and ~53% in Tibet. It was further noted that above the threshold value, the number of co-mutations with low *C_ij_* values (⩽ 0.2) were less while co-mutations with high *C_ij_* (> 0.2) remained unaffected. This observation signifies the role of *p*-value in determining the significant co-mutations pairs. Further, the distribution of *C_ij_* in Andes population showed a heterogeneous distribution of the variations within the samples, i.e, there exist a fewer co-mutations for *C_ij_* > 0.8, indicating minor alleles in question are not always present in the same samples. The number of nodes and edges for all the three regions are also mentioned in Table 1. To get an overall idea for these three regions, we scanned for the common nodes and the common connections among these three co-mutation networks (Fig 2a,b).

**Table 1.** Co-mutation networks V represents the number of nodes, L represents the number of connections in largest connected component, *C_ij_* is the co-mutation frequency, we considered those pairs where *C_ij_* > 0, *p*-value is the measure of statistical significance. We have performed 10,000 random permutations to generate the *p*-value and considered only those pairs with *p*-value ⩽0.05.

| Regions | Sample size | V | L |
| --- | --- | --- | --- |
| Tibet | 86 | 398 | 3459 |
| Ethiopia | 119 | 838 | 13770 |
| Andes | 496 | 1197 | 20224 |

**Table 2.** Global properties of co-mutation networks

| Network property ↓ | Tibet | Ethiopia | Andes |
| --- | --- | --- | --- |
| Average degree | 17 | 33 | 34 |
| Clustering coefficient | 0.8 | 0.7 | 0.8 |
| Modularity | 0.7 | 0.5 | 0.4 |
| Avg path length | 4.3 | 2.8 | 2.3 |
| Degree-degree correlation | 0.4 | 0.2 | -0.4 |

**Fig. 2.**
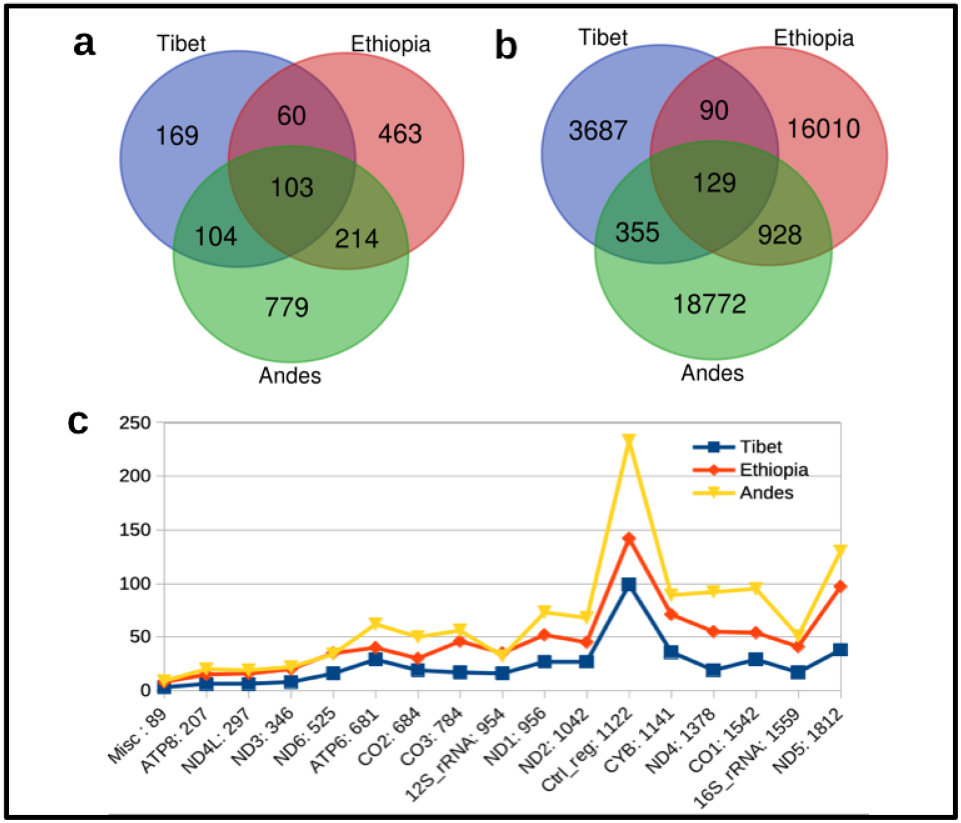
Distribution of nodes **(a)** and co-mutation pairs **(b)** across all the three regions. **(c)** Nodes which are participating in network construction were mapped to their respective genes and genes were counted and plotted on y-axis with their lengths on x-axis. Note that t-RNA genes are not shown.

### 3.2 Structural Properties of Co-mutation Networks

The nodes in these co-mutation networks consist of nucleotide positions which vary among the mitochondrial genome samples for each region. In the co-mutation networks, the degree provides information on the frequency of co-mutation of any given variable site with the other variable sites. A node with high degree corresponds to a variable site undergoing high number of co-mutation with many variable sites. The nodes with the highest degree in the Tibetan (12308, tRNA-Leu) and Andes (10398, ND3) co-mutation networks were commonly present in all the three networks while that of the Ethiopian co-mutation network, the highest degree node (4104, ND1) was present in Andean and was absent in Tibetan network. It is noteworthy that these highest degree nodes were found to be haplogroup markers such that 12308 in Tibetan for K and U haplogroups of N lineage, 4104 in Ethiopian for L0, L1, L2 and L5 haplogroups of L lineage, and 10398 in Andean for multiple haplogroups of L, M and N lineages. The revelation of these haplogroup markers as high degree nodes suggested the dominance of specific haplogroup background for the co-mutation of mtDNA variable sites for each region. Among the high degree nodes, A15301G (Cyb gene) node was found to be common among all the three regions. The corresponding variable site characterizes lineage L and N, and was suggested to be a candidate site for functional analyses and data association [36]. Further, all the three networks were found to have small-world property characterized by high clustering coefficient < *C* >_*real*_ / < *C* >_*rand*_ >> 1 and small diameter *L_real_*/*L_rand_* ~ 1 as for many other real-world networks [38–40]. The small-world behaviour shown by the brain networks suggests swift flow of information in minimal steps from one region to another. Similarly, in co-mutation networks the information of change in allele frequency of a certain nucleotide at one position is transferred to another nucleotide at other position in the same mtDNA sample. Although, for these co-mutation networks, it is a subject of further investigation that whether the two nodes which are connected with each other through more than one step are also sharing the information of change in allele frequency or not. In terms of co-mutation this provides evidence for the fixation and inheritance of variations as a single cohort in the human mitochondrial genome.

Similar to various biological networks [41], a high value of average clustering coefficient (〈*C*〉) was observed here as well for all the co-mutation networks. This depicts high interactions (co-mutations) among the neighbors. Further, a decrease in the tendency of a variable site to form clusters with an increase in the degree of that node (Fig. 3) implies the presence of hierarchy in the network [42, 43]. mtDNA has acquired several variations since humans have first migrated outside Africa throughout the world [44]. Different variations have been enriched in various sub-populations at separate times giving rise to various haplogroups. This temporal and spatial co-evolution of variable sites in mtDNA becomes evident with the presence of high clustering and hierarchy in these co-mutation networks.

**Fig. 3.**
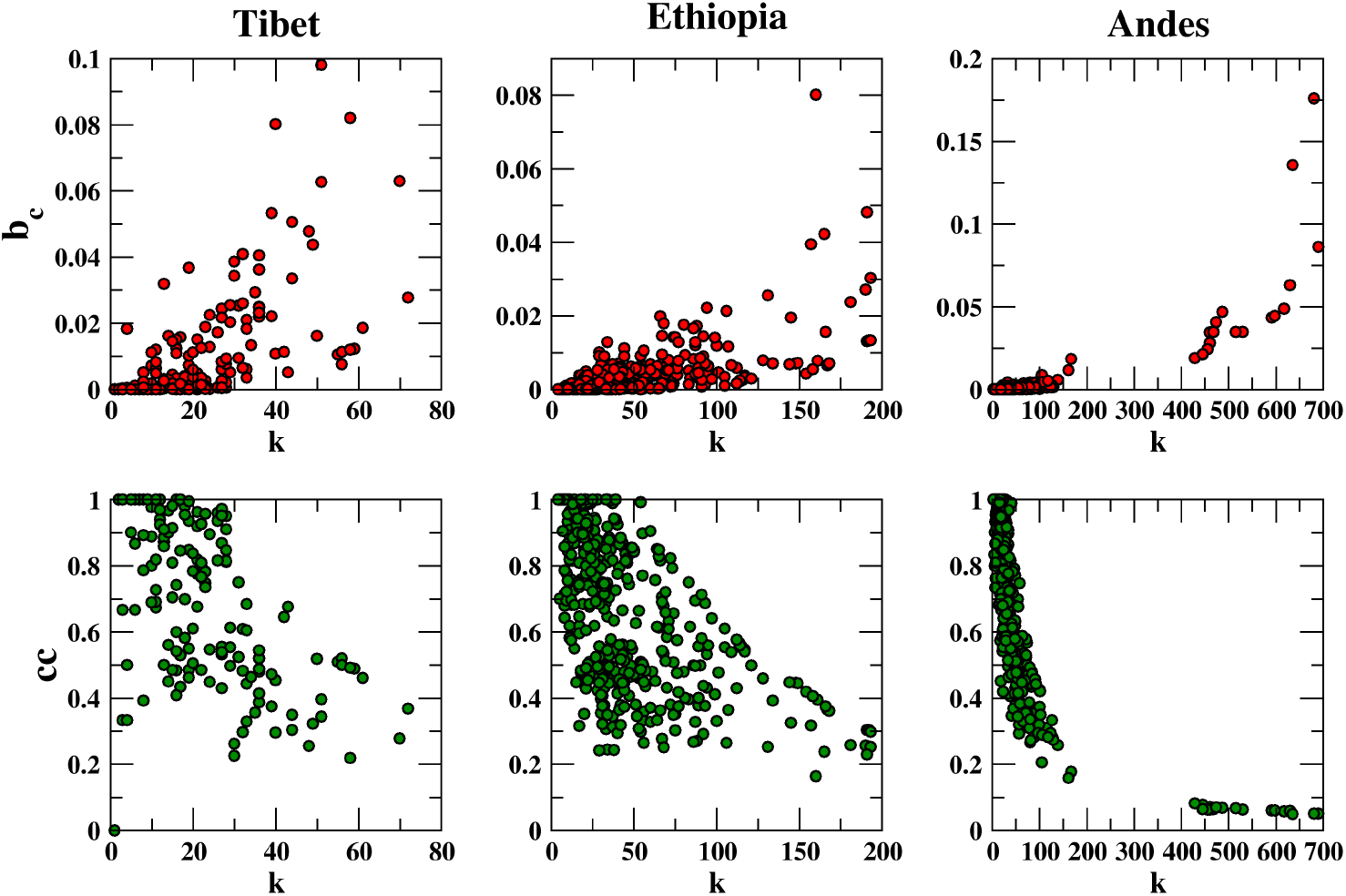
Betweenness centrality and clustering coefficient are plotted as function of degree for all the three regions.

**Fig. 4.**
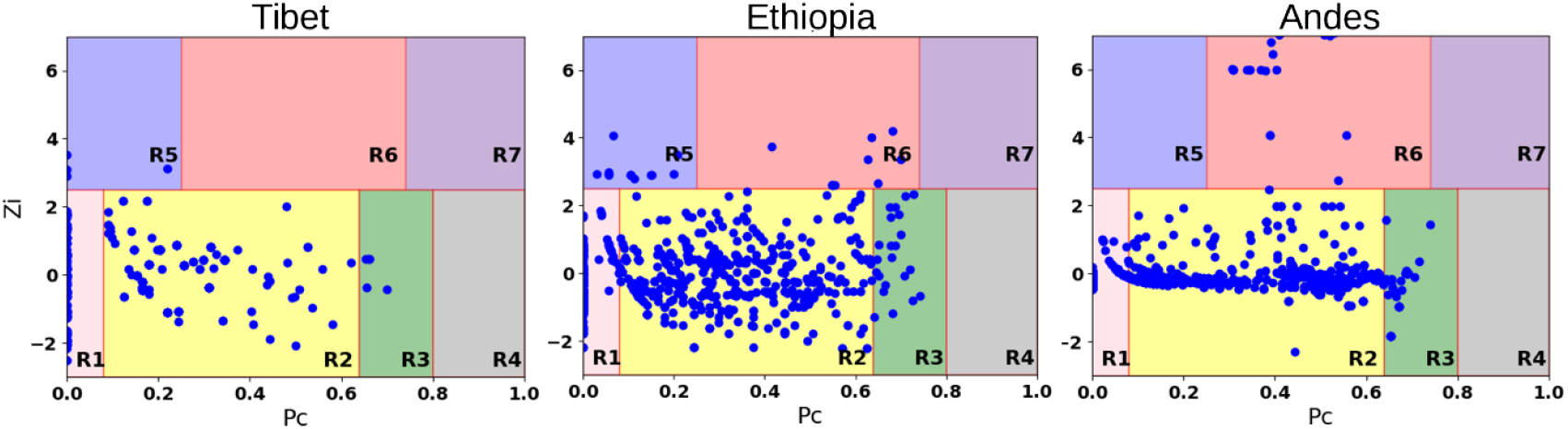
Roles of nodes in Z–P parameter space.

For a network, resilience is an important property, which is measured by the betweenness centrality. Usually, the nodes with high degree tend to have high betweenness centrality. However, it is observed here that a few nodes in spite of having low degree have high betweenness centrality. Which means that these nodes co-mutate with a few variable sites but from different modules. Thus, these sites are significantly important since their removal can result in breakdown of the network. We found four such nodes in the Tibetan co-mutation network which were 709, 15927, 16172 and 16362. Interestingly, all these nodes were again found to be haplogroup markers. This suggests that haplogroup markers not only provide necessary evolutionary background but also play a key role in assisting co-evolution of different clusters. Moreover, 15927 node belongs to tRNA-Threonine which is one of the highly mutated tRNA among all the tRNAs in humans[45], and 709 belong to the 12S-rRNA gene. tRNAs and rRNAs play a central role in protein synthesis. This signifies that tRNAs and rRNAs might play decisive roles in the co-evolution of different mutational cohorts in Tibetan population. On the contrary, we did not find any such nodes in Ethiopian or Andean co-mutation networks. In these two networks the nodes with high betweenness centrality were those which have a large degree. This suggests that in Ethiopian and Andean populations the mtDNA has evolved through continuous co-evolution of different nodes (nodes with high *β_c_* and *k*) of various cohorts while in Tibetan population the mtDNA has evolved through discrete cohorts of co-evolving variable sites (nodes with high *β_c_* and low *k*). This supports the evidence of inheritance of multiple ancestral gene-pools from pre-LGM and post-LGM settlers in the current Tibetan population [46].

We applied Louvain community detection algorithm [34] on co-mutation networks which was implemented in python community module. The high modularity indicates formation of mutational cohorts pertaining to evolutionary constrains at whole genome level. We analyzed the genetic background of the communities formed in these co-mutation networks. On considering only the coding regions, it was observed that one or a few genes predominate each community in all the three co-mutation networks (Supplementary Table 1). Particularly, in Tibet region, *ATP6, CYB, ND5* genes predominated while in Ethiopia *ND5* gene, and in Andes regions *ND5*, and *CYB* genes predominated among all the communities. *CO1* and *ND2* genes were also found to dominate at least in one community in each of the three regions. We also analysed the highest degree nodes for the coding region in each community. These high degree nodes are considered as “community cores”. We found that none of these community cores are common among the three regions. Although the three regions have certain number of common nodes (Fig 2), the community structures are derived by independent nodes. Upon individual inspection of the communities we found that in each of the three regions despite contributing few nodes *tRNA-Leu, tRNA-Lys, and tRNA-Gln* were found to be community cores in the Tibet, Ethiopia and Andes regions, respectively. Apart from that, the *CR* was evenly present in all the communities. We investigated localization properties of eigenvectors of these co-mutation networks. Localization of eigenvectors enjoy wide range of application in networks; in disease spreading[52], perturbation of propagation in mutualistics networks[53]. Other application of localization can also be found in [55, 56]. To quantify localization, we use correlation dimension (*D*_2_) calculated by using box counting method for multifractal analysis of eigenvectors[56]. If *D*_2_ →0, eigenvector is said to be localized while if *D*_2_ →1, eigenvector is considerd as delocalized. Thus, *D*_2_ provides insight about the degree of localization of eigenvectors. We focus on the eigenvectors of eigenvalues nearer to zero and *D*_2_ is average over all the eigenvectors inside width *dλ* = 0.5. Note that, slight increase or decrease in the width will not alter the results. Tibetan and Andean networks were found to be more localized with *D*_2_ being 0.43 compared to Ethiopian network with *D*_2_ 0.65. In these networks the tendency of comutation is captured by *D*_2_ in terms of localization. A co-mutation occurs when two minor alleles are present in considerable frequency in the population and the tendency of further co-mutation is limited by introduction of new alleles. In Andean and Tibetan populations this tendency have been limited compared to Ethiopia, which might be possible due to recent population admixture experienced by Ethiopian population[57]. For these networks, there exists no node with its degree distinctly very high than those of the other nodes, and hence importance of a node can not assigned based on its degree. Nevertheless, due to the presence of the high modularity, importance of a node can be determined, to a great extent, by its within-module degree and participation coefficient, which defines how a node is positioned in its own module and with respect to other modules [47–49]. Based on the within-module degree and the participation coefficient, nodes were categorized as module hubs and non-hubs. The highest degree nodes for the Tibetan co-mutation network were found in R3, non-hub connector category, while for the Ethiopian and Andean co-mutation networks the highest degree nodes were found in R6 connector hubs category. In the Tibetan co-mutation network, the nodes in R5 category were 3010 (*16S_rRNA*), 8414 (*ATP8*), 14668 (*ND6*) and 12361 (ND5). Variable site 3010 was shown to be a high-altitude marker in Tibetan population [50] and also form co-occurrence motifs with variable sites 8414 and 14668[31] while, 12361 was shown to be associated with nonalcoholic fatty liver disease [51]. It is noteworthy that in the Tibetan co-mutation network there were no nodes in R6 category, and in the Andean co-mutation network there were no nodes in R5 category while in the Ethiopian co-mutation network nodes were present in both the R5 and R6 categories. These co-mutation networks are then analysed at gene level through the construction of the gene-gene interaction networks discussed further.

### 3.3 Gene-Gene Interaction (GGI) Networks

First, we mapped the nodes to their corresponding genes and counted the occurrence of variable sites pertaining to each gene to access these occurrences with respect to their lengths for each region (Fig. 2c). It is reflected from the figure that except the *Control region (CR)*, the occurrence of variable sites for each gene increases with an increase in the length of genes. It is quite evident that certain genes are contributing more to the variable sites in the network construction than others in a particular region. Especially, *ATP6, CO2* and *ND2* genes are contributing equally in the Tibet and Ethiopia networks while *12S-rRNA, 16S-rRNA, CO3* and *ND4* are contributing equally in the Tibet and Andes. Among the coding genes, ND5 gene showed highest difference of contributing nodes with minimum in Tibet and maximum in Andes (Fig. 2c). The contribution of each gene per 100 samples for the network construction was highest in Ethiopia among all the regions. The nodes pertaining to *CR* displayed largest participation in the network construction due to the fact that it is the highly variable part of mtDNA [35]. Contribution of the variable sites in each gene yields a partial information about the interaction of the genes. To overcome this limitation, we generated the gene-gene interaction networks through mapping the variable sites with the respective genes as discussed in the Methods section.

The gene-gene interaction networks provide a holistic and reductionist approach to investigate interactions in the three high altitude regions. We found significantly 17 gene-pairs in Tibetan, 23 gene-pairs in Ethiopian and 44 gene-pairs in Andes population after comparing with their corresponding random networks. Among these, the pair with the highest weight in the Tibetan network was *ATP6-CYB*, in the Ethiopian it was *ND4-ND5* and in the Andes it was *CYB-ND4*. Four significant gene-gene pairs were found to be commonly present in all the three populations, which were *CO1-CO2, ND2-ND4, ND3-ND4* and *ND4-ND5*. All these genes are involved in oxidative phosphorylation pathway (KEGG entry: 00190) and thermogenesis pathway (KEGG entry: 04714) [58], along with that these genes are also found to be involved in cellular respiration (GO:0045333) and response to abiotic stimulus (GO:0009628) [59]. Since the three populations are believed to share a similar physical environment and undergoing the process of convergent evolution, we analyzed the properties of all the common nodes (variable sites) and extracted their co-mutations information and mapped these co-mutations to corresponding genes for each population to obtain co-evolution network of genes. The common variable sites categorize these three populations under the same haplogroups while the co-mutations of the involved nodes apparently differ in these three populations. To capture this difference at the genetic level, significant genetic interaction (Fig 5) of the common nodes were extracted based on the functional enrichment analysis for GO terms and KEGG pathways using DAVID [60]. It was found that in the Tibetan population *CO3, CYB* and *ND5* genes, in the Andean population *ATP6, CO3, CYB, ND3* and *ND4* genes, and in the Ethiopian population *ATP6, CO1, CO2, ND1, ND2, ND4* and *ND5* genes were significantly interacting with other genes. The functional enrichment of these genes sets is shown in table in Fig 5. It is noteworthy that from cytochrome oxidase complex, the *CO3* sub-unit was interacting in the Tibetan and Andean populations while *CO1* and *CO2* sub-units were interacting in Ethiopian population. *CO3* sub-unit is the putative site for the entry of oxygen into the large cytochrome oxidase complex thereby regulating its activity under hypoxic conditions [61]. Even though, Ethiopian gene set has not shown any feature related to the hypoxia adaptation in functional enrichment analysis, variations in *ND1* and *ND2* genes were shown to be associated with high-altitude hypoxia in Tibetan yak [62] and endemic Ethiopian rats [63]. Apart from the functional enrichment, the Tibetan and Andean gene-sets were also involved in non-alcoholic fatty liver disease (NAFLD). However, it has been shown that high altitude might improve the mitochondrion function and alleviate the NAFLD [64]. Further to explore the role of such genetic interactions at region specific level, we extracted the co-mutation pairs pertaining to the exclusive nodes of each region and constructed GGI networks. In GGI networks with these exclusive nodes, we found that there are certain interactions which were having significantly lower weight and certain others with significantly higher weight than the corresponding random ones (Fig. 6). It is readily observed from Fig. 6 that those genetic interactions which were significantly up in Andean and Tibetan population were significantly down in the Ethiopian population and vice versa.

**Fig. 5.**
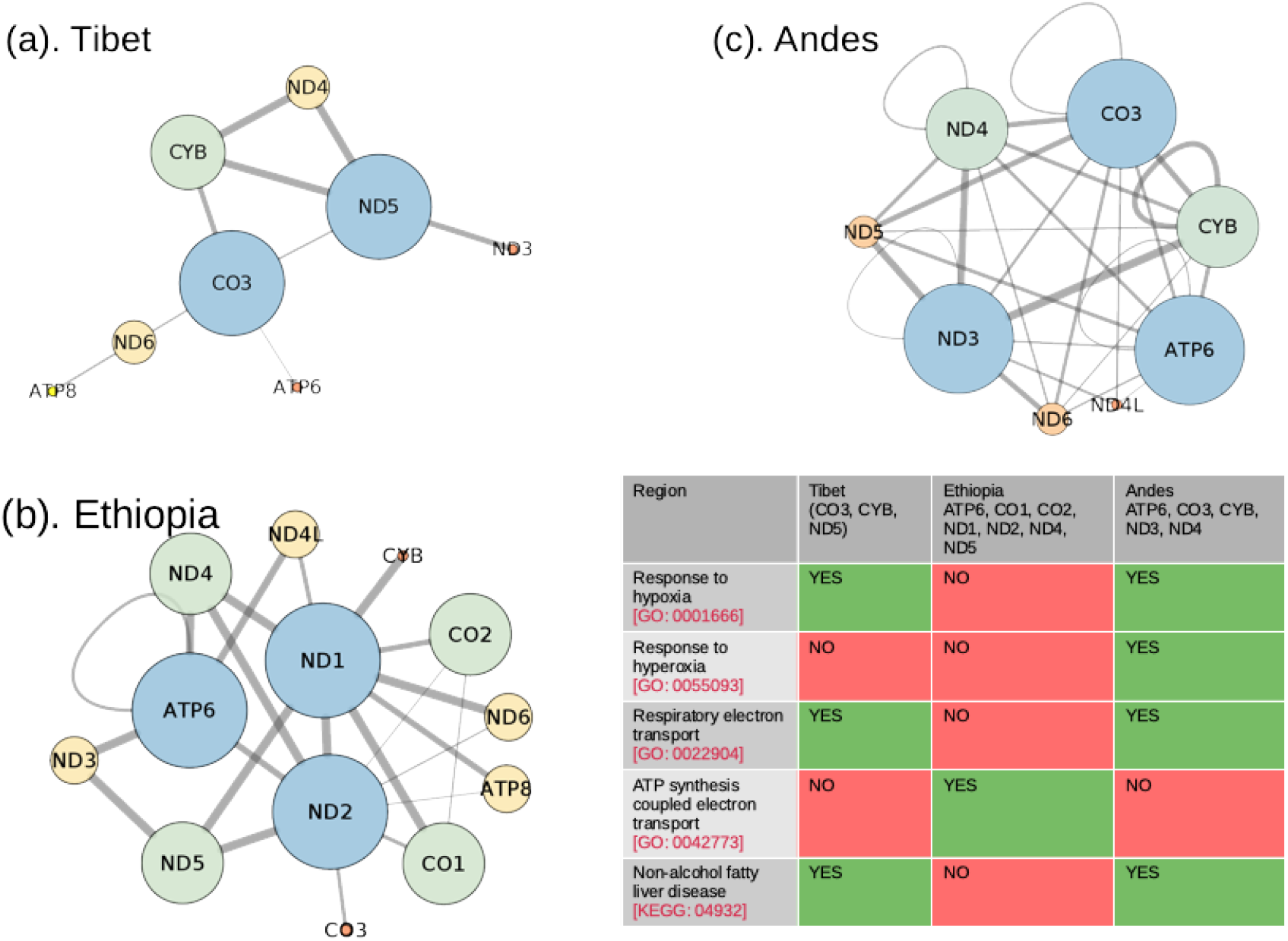
Significant gene-gene interactions of common node co-mutations and corresponding GO terms and KEGG pathway. (The node size depicts the degree of the node and edge size represents the edge weight.)

**Fig. 6.**
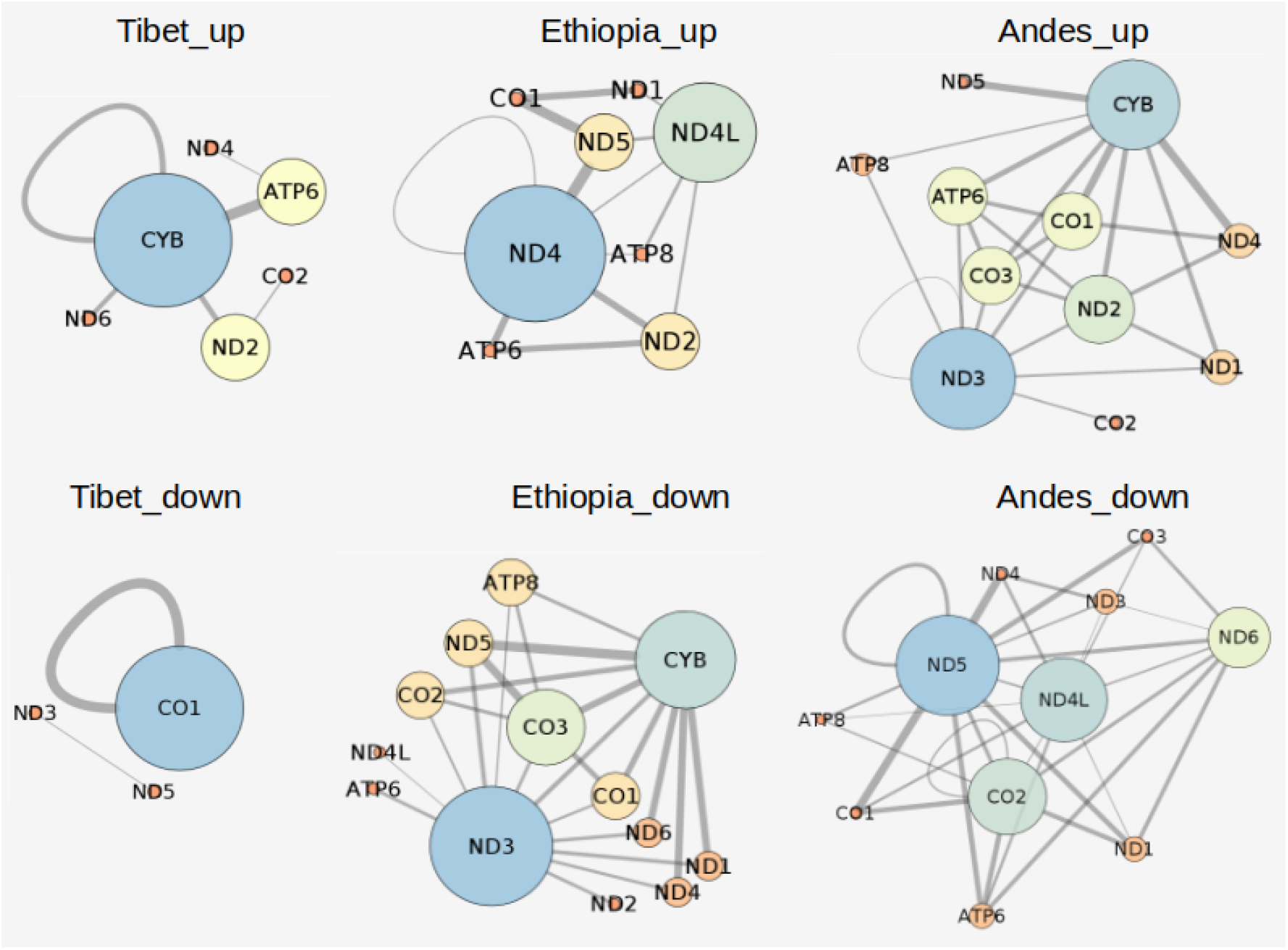
Significant gene-gene interactions of exclusive nodes co-mutations. The *Region_up* and *Region_down* represents those interactions which were showing higher and lower weight than the corresponding random networks for each region, respectively. (The node size depicts the degree of the node and edge size represents the edge weight.)

This suggests that both Tibetan and Andean populations have evolved at high altitudes through the interactions of *CYB* and *CO3* genes while Ethiopian population deviated in sharing the mitochondrial genetic interactions with the other two populations. This dissimilarity could be explained based on two facts, one is that Ethiopia is situated at lowest altitude among all the three populations, and second that Ethiopia is believed to undergo frequent admixture in its gene pool from lower altitude populations[65].

## 4. Conclusion

Although it is true that biologically a population can be defined in terms of haplogroups or specific mutations, with a firmly significant threshold co-mutations fortify the specific genetic interactions even in similar environmental backgrounds. As our result showed that in Andean and Tibetan populations, mtDNA has evolved similarly for these populations while Ethiopian population showed different signatures, suggesting that Ethiopian population has evolved at high-altitudes through some different mechanisms than Tibetan and Andean populations. In Tibetan and Andean population, *CYB* and *CO3* genes are commonly present in Tibetan and Andean population and are interacting with other genes namely *ND5* in Tibetan and *ATP6, ND3* and *ND4* in Andean. Whereas, in Ethiopian population four NADH dehydrogenase genes (*ND1, ND2, ND4, ND5*) showed interactions with *ATP6, CO1* and *CO2*. It is noticeable that in exclusive GGI networks, Ethiopian population showed the contrasting behaviour compared to the Tibetan and Andean population at the genetic level. Further, the *D*_2_ analysis also showed that Tibetan and Andean populations are similar in their localization behavior compared to Ethiopian population. Comprehensively, our analysis showed that Ethiopian population has evolved heterogeneously than the Tibetan and Andean populations. The method of co-mutation analysis provides information about coevolving sets of genes based on their co-mutations to further investigate their role in environmental adaptations pertaining to convergent evolution at high-altitudes in human sub-populations.

## Acknowledgments

SJ acknowledges support from SERB grant (EMR/2016/001921) and the council of scientific and industrial research grant (CSIR, 25(0293)/18/EMR-II) Govt of India. RKV gratefully acknowledges University Grant Commission for SRF fellowship (305089). MI acknowledges the Ministry of education and science of the Russian Federation megagrant (075-15-2019-871). We acknowledge Pramod Shinde for continuous discussions throughout and Cristina for useful discussions in the beginning of the project.

## Notes

### Competing Interest Statement

The authors have declared no competing interest.

